# Nociceptors use multiple neurotransmitters to drive pain

**DOI:** 10.1101/2025.09.09.675093

**Authors:** Donald Iain MacDonald, Rakshita Balaji, Alexander T. Chesler

**Author notes:** Corresponding author:* (DIM) or (ATC).

## Abstract

Nociceptors are excitatory neurons that express a range of neuropeptides and have the essential role of detecting noxious mechanical, thermal and chemical stimuli. Ablating these neurons profoundly reduces responses to pain. Here we investigated how nociceptive information is transmitted by developing genetic approaches to suppress glutamate transmission and neuropeptide signaling, individually and in combination. Remarkably, many pain responses persisted in mice where either nociceptor glutamate or neuropeptide signaling was blocked. By contrast, mice lacking both glutamate and neuropeptide transmission in nociceptors displayed profound pain insensitivity closely matching the effects of cell ablation. Together our results establish a role for neuropeptides as *bone fide* pain transmitters and demonstrate redundancy in nociceptor signaling, resolving long-standing questions about how pain is communicated to the brain.

---

People born without functional nociceptors often succumb to injuries and die young, illustrating the crucial role of pain sensation (*1*). However, ongoing activity in nociceptors can also trigger maladaptive chronic pain, a leading cause of disability worldwide (*2*). Over the past 30 years, the discovery of TRP channels and voltage-gated sodium channels specifically expressed in nociceptors has produced a detailed picture of how tissue damage is converted into nociceptor action potential firing (*3*–*5*), and spurred the development of a new class of painkillers (*6*). By contrast, how nociceptor activity is translated into neurotransmitter release to communicate with the spinal cord and brain to drive pain remains poorly defined (*7, 8*).

Nociceptors release multiple different signaling molecules from both their central and peripheral terminals, many of which have been implicated in driving pain (*9*–*11*). While glutamate is thought to be the major excitatory neurotransmitter (*12*–*14*), nociceptors also co-release a cocktail of neuropeptides, including Substance P (SP), Calcitonin gene-related peptide (CGRP), Neurokinin A (NKA), Neuromedin B (NMB), Galanin, and Pituitary Adenylate Cyclase Activating Polypeptide (PACAP) (fig. S1) (*15*). Classic studies suggested Substance P was the principal neurotransmitter released by C fibres (*16*), and CGRP is a key mediator of pain in migraine (*17*). However, deletion of Substance P, CGRP or both in combination has little to no effect on acute pain behavior in mice (*18*–*20*). Electron microscopy studies have revealed that multiple neuropeptides are contained within the same dense core vesicle (*21*), while the corresponding post-synaptic receptors are often co-expressed (*22*), consistent with possible functional overlap (*23*).

Here, through systematic analysis of animals lacking different combinations of neuropeptides and glutamate signaling, we uncovered a striking functional redundancy at the level of nociceptor neurotransmission. Our findings challenge longstanding assumptions about the organizational logic of ascending pain pathways and suggest that nociceptors use multiple parallel pathways to ensure the robust and reliable transmission of pain.

## Results

### Loss of glutamate transmission from nociceptors does not abolish pain

The capsaicin receptor Trpv1 is developmentally expressed in the great majority of nociceptors (*24*). As reported previously (*25*), we found that diphtheria toxin ablation of Trpv1 lineage nociceptors resulted in dramatic insensitivity to heat, cold and chemical pain stimuli (Figure 1A-F). We next generated mice lacking vesicular glutamate transporter 2 (VGLUT2) in Trpv1 lineage nociceptors and directly compared their ability to sense pain with the nociceptor-ablated mice. In line with prior studies (*12*–*14*), the VGLUT2 cKO^Trpv1^ mice developed self-inflicted injuries and exhibited decreased sensitivity to heat (Fig. 1A-C). However, the animals lacking VGLUT2 were much more sensitive to heat than the mice where Trpv1 neurons were ablated. The VGLUT2 cKO^Trpv1^ mice also showed a surprisingly normal response to cold (Fig. 1D), and a vigorous, albeit delayed, response to the chemical algogens capsaicin and mustard oil (Fig. 1E-F, fig. S2). Thus, suppressing glutamate transmission resulted in measurable but modest pain deficits when compared to the profound analgesia produced by ablating all Trpv1 lineage neurons. Importantly, VGLUT2 deletion does not trigger compensatory increases in expression of other vesicular glutamate transporter genes (*13*). Taken together, these experiments suggest that nociceptors must use other neurotransmitters to transmit pain.

**Figure 1.**
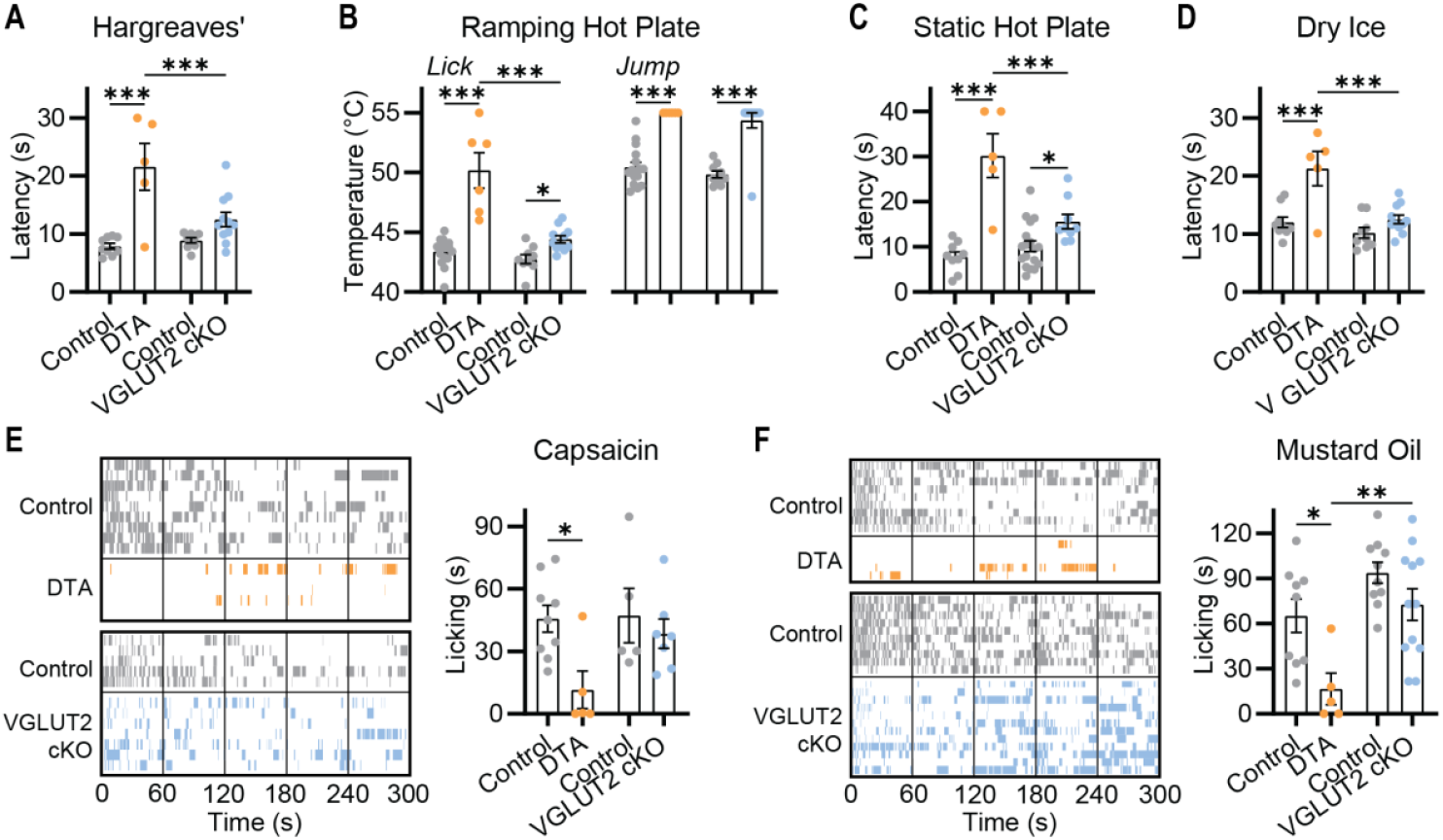
Deletion of VGLUT2 does not recapitulate the loss of pain seen in nociceptor-ablated mice. **(A-F)**. Pain behavioral assessment of Trpv1-Cre DTA and Trpv1-Cre VGLUT2 cKO mice and their respective littermate controls. **(A)** Latency to withdrawal to radiant heat. n=9 (5M/4F) for Control and n=5 (4M/1F) for DTA; n=9 (4M/5F) for Control and n=11 (6M/5F) for VGLUT2 cKO. **(B)** Temperature of first lick and jump on Ramping Hot Plate. n=16 (5M/11F) for Control and n=6 (4M/2F) for DTA; n=9 (4M/5F) for Control and n=11 (6M/5F) for VGLUT2 cKO. **(C)** Latency to withdrawal on Static Hot Plate. n=9 (5M/4F) for Control and n=5 (4M/1F) for DTA; n=18 (11M/7F) for Control and n=9 (6M/3F) for VGLUT2 cKO. **(D)** Latency to withdrawal to dry ice. n=9 (5M/4F) for Control and n=5 (4M/1F) for DTA; n=9 (4M/5F) for Control and n=11 (6M/5F) for VGLUT2 cKO. **(E)** Ethogram and summary data showing time spent licking paw injected with capsaicin. n=9 (5M/4F) for Control and n=5 (4M/1F) for DTA; n=5 (3M/2F) for Control and n=7 (5M/2F) for VGLUT2 cKO. **(F)** Ethogram and summary data showing time spent licking paw injected with mustard oil. n=9 (5M/4F) for Control and n=5 (4M/1F) for DTA; n=10 (3M/7F) for Control and n=12 (5M/7F) for VGLUT2 cKO. Bars represent mean ± SEM. Means were compared by One-WAY ANOVA, with Holm-Sidak multiple comparisons test used to compare DTA and VGLUT2 cKO with one another, and with their respective littermate controls. \**p*<0.05, \*\**p*<0.01 \*\*\**p*<0.001.

### The enzyme PAM is essential for neuropeptide signaling from nociceptors

The persistence of many nociceptive behaviors in the absence of glutamate release implicates neuropeptides as potential transmitters of pain (Fig. 2A). However, constitutive knockout of specific peptides has little effect on pain sensitivity (*18*–*20*). Therefore, we implemented a platform to generally suppress neuropeptide signaling. The enzyme peptidylglycine α-amidating monooxygenase (PAM) directs C-terminal amidation of many neuropeptides (Fig. 2B) (*26*). Since this is the final step in peptide biosynthesis, conditional deletion of PAM may provide a strategy to explore the role of neuropeptide signaling in pain (*27, 28*). To validate the efficacy of this approach, we deleted PAM in all caudal peripheral somatosensory sensory neurons by crossing PAM^fl/fl^ mice with animals expressing Hoxb8-Cre (*29*). These PAM conditional knockout (cKO) mice were indistinguishable from their control littermates but, as expected, exhibited no PAM expression in the spinal cord (fig. S3A).

**Figure 2.**
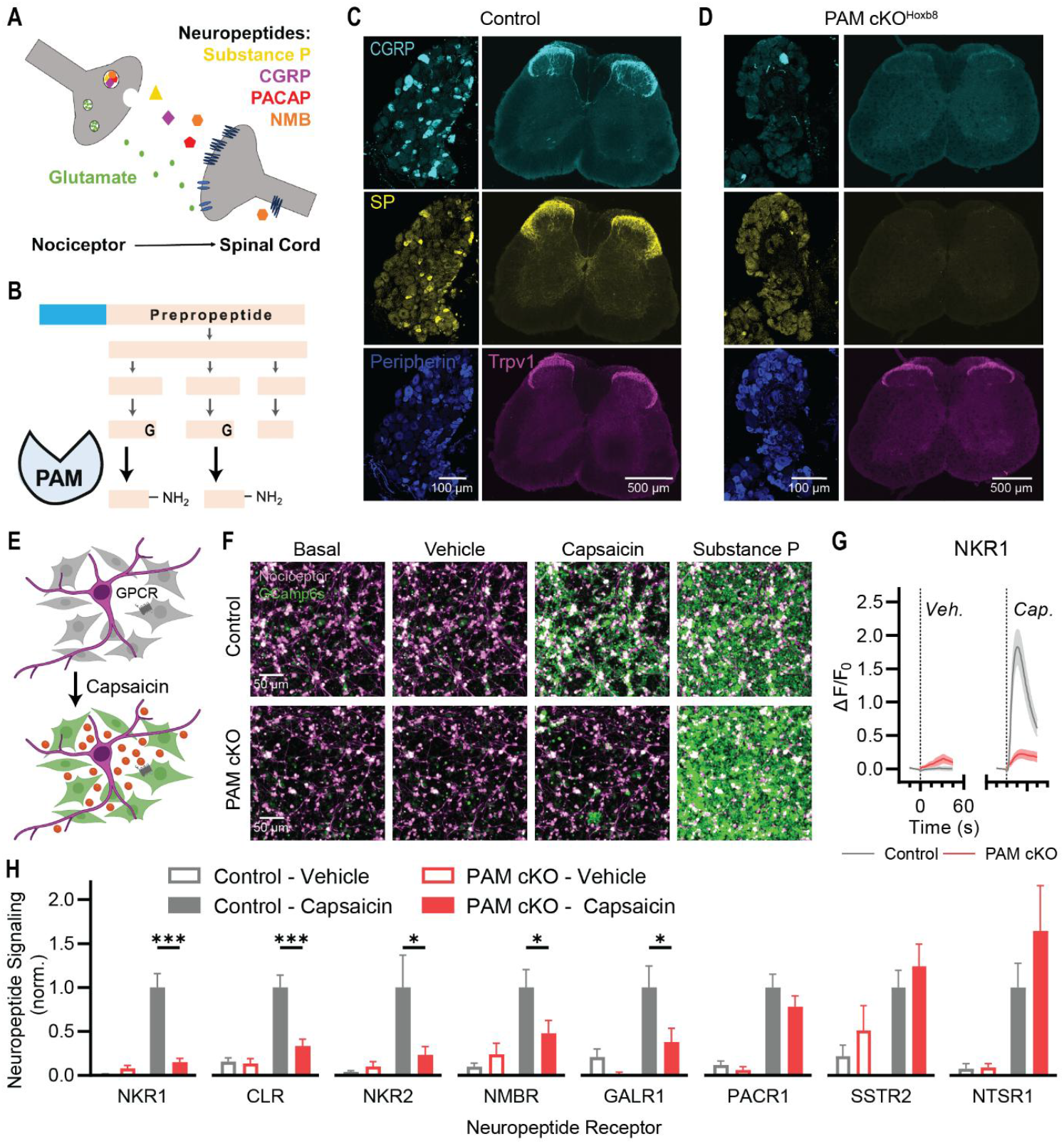
Deletion of PAM suppresses amidated neuropeptide signaling from nociceptors. **(A)** Nociceptors release multiple neuropeptides. **(B)** The enzyme PAM directs the C-terminal amidation of many of these nociceptor peptides. **(C)** Control mice show strong staining for the neuropeptides Substance P and CGRP in dorsal root ganglia and spinal cord dorsal horn. **(D)** PAM cKO_Hoxb8_ show a near complete loss of staining for mature Substance P and CGRP. **(E)** A cell-based biosensor assay enables detection of neuropeptides via their native receptors. **(F)** Capsaicin evokes Substance P release and NKR1 activity from control nociceptors, but not from PAM cKO_Avil_ nociceptors. **(G)** Quantification showing the effect of vehicle and capsaicin on biosensor fluorescence in co-culture with control and PAM cKO nociceptors. **(H)** Quantification of capsaicin-evoked neuropeptide signaling from control and PAM cKO_Avil_ nociceptors using a range of receptor biosensors for different peptides. PAM deletion inhibits SP, CGRP, NKA, NMB, and Galanin signaling. PACAP, NTS and SST signaling are unaffected. NKR1: n=17 for Control; n=15 for PAM cKO. CLR: n=23 for Control; n=20 for PAM cKO. NKR2: n=13 for Control; n=12 for PAM cKO. NMBR: n=24 for Control; n=24 for PAM cKO. GALR1: n=16 for Control; n=17 for PAM cKO. PACR1: n=21 for Control; n=16 for PAM cKO. SSTR2: n=22 for Control; n=21 for PAM cKO. NTSR1: n=15 for Control; n=15 for PAM cKO. All data from at least three independent cultures. Bars represent mean ± SEM. For each biosensor, means were compared using 2-way ANOVA followed by post-hoc Holm-Sidak’s multiple comparison’s test. \**p*<0.05, \*\**p*<0.01 \*\*\**p*<0.001.

Substance P and CGRPα, the major neuropeptides in nociceptors, are C-terminally amidated and were readily detected in nociceptor cell bodies and their central terminals in the spinal cord (Fig. 2C). Notably, the somata and terminals of PAM cKO^Hoxb8^ mice were devoid of this staining (Fig. 2D). By contrast, expression of Trpv1 (Fig. 2D), Peripherin (Fig. 2D) and VGLUT2 (fig. S3B) was unaffected by PAM deletion, indicating nociceptors developed normally in the absence of amidated neuropeptides.

Next, we generated sensory neuron-specific cKO mice using Avil-Cre *(30)* and measured the effects of PAM deletion at a functional level *(31)*. Control and PAM cKO^Avil^ sensory neurons were co-cultured together with HEK293 cells expressing neuropeptide G-protein-coupled-receptors, human Gα15 and GCaMP, enabling us to quantify receptor activation downstream of capsaicin-induced neuropeptide release from nociceptors (Fig. 2E) *(32)*. Stimulating nociceptors lacking PAM evoked minimal activity of biosensor cells expressing the Substance P or CGRP receptors in comparison with the robust activation seen in controls (Fig. 2F, G and H). Indeed, this loss of activity recapitulates the effect of Tac1 and Calca knockout (18), confirming PAM is essential for Substance P and CGRP signaling and validating our approach.

Nociceptors express numerous neuropeptides that show distinct patterns of expression across neuronal subtypes (*15*) (fig. S1) and which, in some cases, are expressed only after injury or in culture (*33, 34*). To comprehensively assess which of these are PAM-dependent, we generated eight new receptor bioassays to detect both unamidated and amidated nociceptor neuropeptides (fig. S4A-L). Dose response curves demonstrate the exquisite sensitivity and selectivity of these lines for their corresponding neuropeptides (fig. S4A-L). As shown in Fig. 2H and fig. S4M, we found that PAM cKO^Avil^ had no effect on capsaicin stimulated release of the unamidated neuropeptides Somatostatin (SST), Neurotensin (NTS) and the endogenous opioid peptides from DRG neurons. By contrast, signaling associated with the amidated peptides NKA, NMB, and Galanin, and Gastrin-release peptide (GRP) was strongly attenuated. The only amidated peptide with normal signaling in the cKO-neurons was PACAP, consistent with previous observations that C-terminal amidation is not required for PACAP to activate its receptor (*35*). Together, these data show that PAM cKO mice lack most amidated neuropeptide function, including the key nociceptor neuropeptides SP, CGRP, NKA, NMB and Galanin.

### Loss of neuropeptide signaling from nociceptors attenuates neurogenic inflammation

The axon flare reflex is thought to involve release of Substance P and CGRP from nociceptor peripheral terminals (*36*). Since both these neuropeptides require C-terminal amidation to signal, we quantified flare responses *in vivo* using laser speckle contrast imaging. Topical mustard oil application to the paw locally increased dermal blood flow in control mice. This flare response was greatly diminished (Fig 3A, B and C) in mice with nociceptor-specific deletion of PAM (PAM cKO^Trpv1^), demonstrating a function for neuropeptide transmission *in vivo*. Interestingly, in DTA^Trpv1^ mice, flare was completely lost (Fig 3D, E and F), insinuating that other mediators are involved in triggering this response.

**Figure 3.**
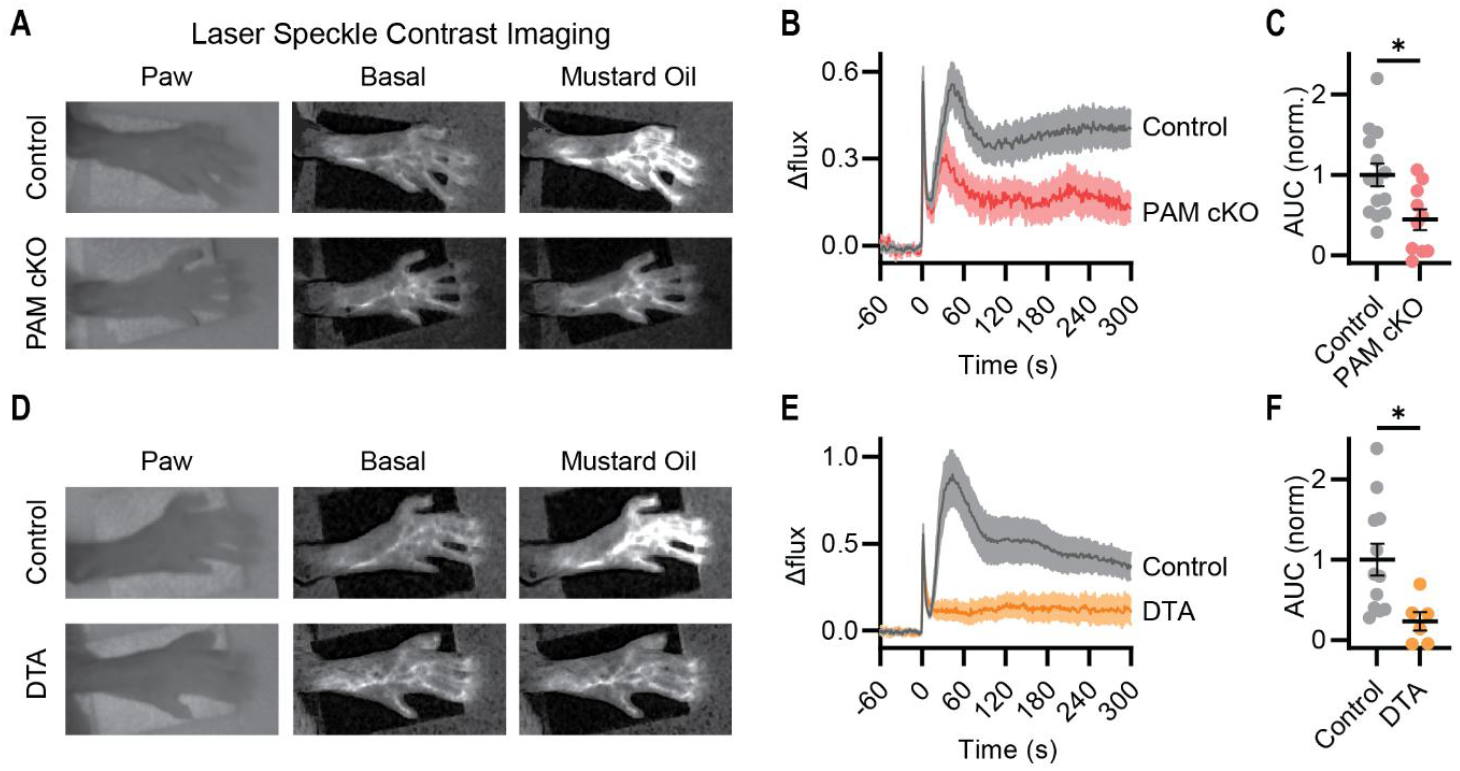
Deletion of PAM in nociceptors inhibits the axon flare response. **(A-F)** Laser speckle contrast imaging was used to measure the blood flow in the paw following nociceptor stimulation with mustard oil. Images **(A)**, time course **(B)** and quantification (area under curve, normalized to control) **(C)** showing that mustard oil application evokes less blood flow in the paw of PAM cKO mice compared to controls. n=14 (5M/9F) for Control and n=10 (7M/3F) for PAM cKO. This response was completely lost in Trpv1-Cre DTA mice **(D-F)**. n=12 (5M/7F) for Control and n=6 (4M/2F) for DTA. Bars represent mean ± SEM. Normalized area under the curve was compared using an unpaired t test. \**p*<0.05, \*\**p*<0.01 \*\*\**p*<0.001.

### Loss of neuropeptide signaling from nociceptors does not abolish pain

Previously we showed that germline global knockout of Substance P and CGRP together had no effect on pain sensitivity (*18*). The PAM cKO^Trpv1^ strategy eliminates the majority of neuropeptide function selectively in nociceptors. Surprisingly the pain behavior of these mice was also largely normal. Reflexive responses to heat and cold were indistinguishable from control littermates (Fig. 4A and D). On Static and Ramping Hot Plates, the mice were moderately less sensitive than littermate controls (Fig. 4B and C), reminiscent of the effects of VGLUT2 deletion in nociceptors (Fig. 1). Responses to capsaicin and mustard oil were also very similar across genotypes (Fig. 4E and F).

**Figure 4.**
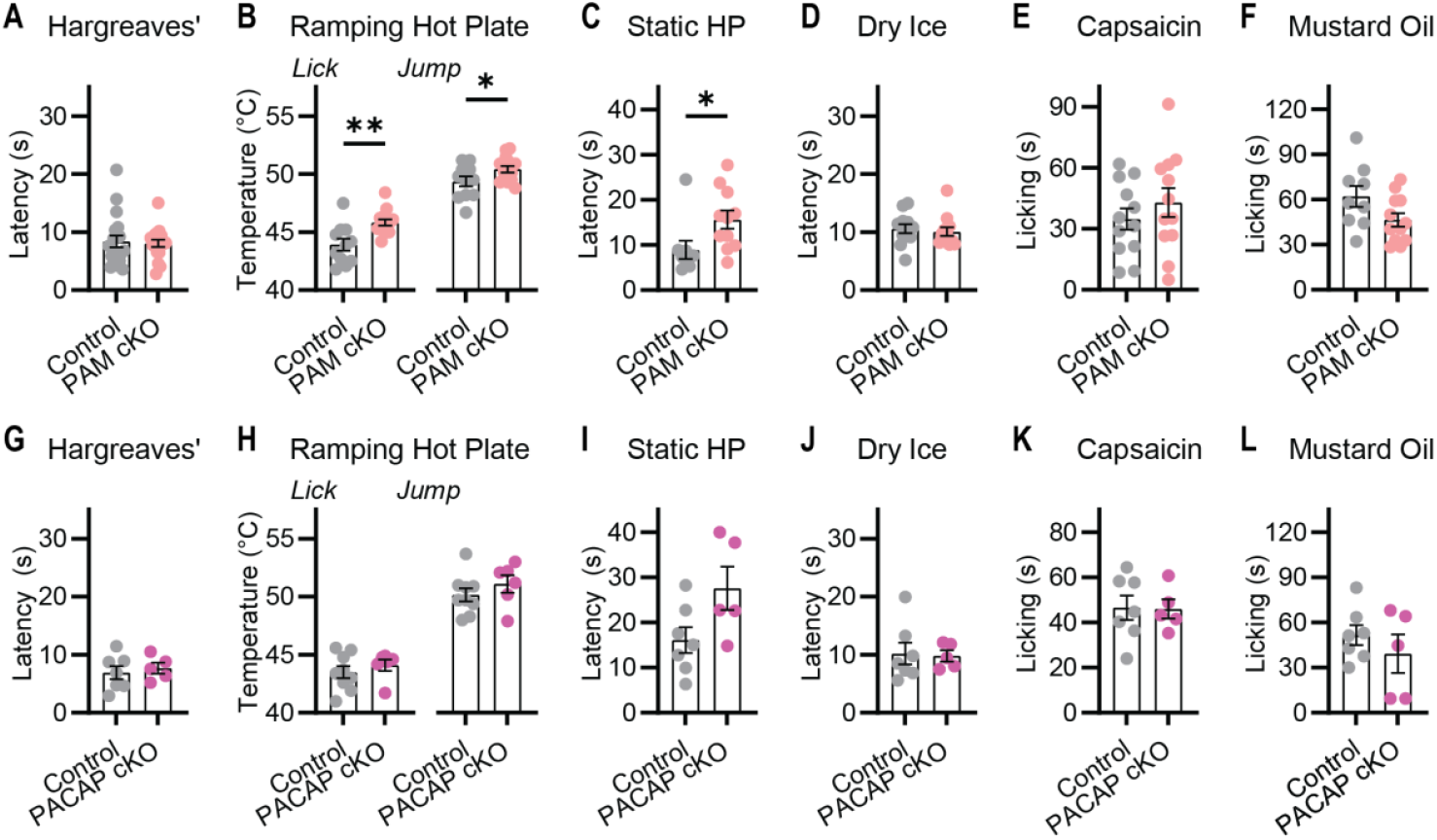
Mice lacking neuropeptides show largely normal pain behavior. **(A-F)** Pain behavioral assessment of PAM cKO mice. **(A)** Latency to withdrawal to radiant heat. n=19 (9M/10F) for Control and n=18 (9M/9F) for PAM cKO. **(B)**Temperature of first lick and jump on Ramping Hot Plate. n=11 (6M/5F) for Control and n=14 (8M/6F) for PAM cKO.**(C)**Latency to withdrawal on Static Hot Plate. n=9 (3M/6F) for Control and n=11 (6M/5F) for PAM cKO. **(D)** Latency to withdrawal to dry ice. n=12 (6M/6F) for Control and n=12 (6M/6F) for PAM cKO. **(E)** Time spent licking paw injected with capsaicin. n=12 (6M/6F) for Control and n=12 (8M/4F) for PAM cKO. **(F)** Time spent licking paw injected with mustard oil. n=9 (2M/7F) for Control and n=12 (3M/9F) for PAM cKO. **(G-L)** Pain behavioral assessment of PACAP cKO mice. **(A)** Latency to withdrawal to radiant heat. n=7 (4M/3F) for Control and n=5 (3M/2F) for PACAP cKO. **(B)** Temperature of first lick and jump on Ramping Hot Plate. n=9 (5M/4F) for Control and n=6 (3M/3F) for PACAP cKO. **(C)** Latency to withdrawal on Static Hot Plate. n=7 (3M/4F) for Control and n=5 (2M/3F) for PACAP cKO. **(D)** Latency to withdrawal to dry ice. n=7 (4M/3F) for Control and n=5 (3M/2F) for PACAP cKO. **(E)** Time spent licking paw injected with capsaicin. n=7 (4M/3F) for Control and n=5 (3M/2F) for PACAP cKO. **(F)** Time spent licking paw injected with mustard oil. n=7 (4M/3F) for Control and n=5 (3M/2F) for PACAP cKO. Bars represent mean ± SEM. Means compared using an unpaired t test. \**p*<0.05, \*\**p*<0.01 \*\*\**p*<0.001.

The minor effects of deleting PAM on pain behavior are unlikely to reflect compensatory changes in gene expression as no significant transcriptomic changes were detected (fig. S6), or the efficiency of PAM cKO^Trpv1^ (fig. S7). We also generated a nociceptor specific knockout of PACAP, the major neuropeptide in nociceptors that functions in the absence of PAM (Fig. 2) and validated loss of PACAP transmission (fig. S8). However, these mice also exhibited normal pain responses (Fig. 4G, H, I, J, K and L).

### Blocking both glutamate and neuropeptide transmission from nociceptors leads to pain insensitivity

Our results reveal that individually blocking either glutamate (Fig. 1) or neuropeptide signaling (Fig. 4) has minimal effects on pain, particularly in comparison with the profound analgesia produced by nociceptor ablation (Fig. 1). This provokes the hypothesis that glutamate and neuropeptides are mutually redundant transmitters of pain. This leads to the clear prediction that simultaneously eliminating both glutamate and neuropeptide signaling should completely block pain transmission. We therefore generated mice lacking both PAM and VGLUT2 in nociceptors (PAM VGLUT cDKO^Trpv1^). Remarkably, these animals showed a profound loss of acute pain equivalent to ablating the Trpv1 lineage (Fig. 5). PAM VGLUT2 cDKO^Trpv1^ animals showed dramatically reduced responses to heat (Fig. 5A, B and C), cold (Fig. 5D), and the injection of chemical algogens (Fig. 3E and F, and fig. S9). Notably, many animals reached experimental cut-off values and the analgesia was far greater than the modest effects of deleting either PAM or VGLUT2 alone. This synergistic phenotype cannot be explained by denervation of the central terminals as Trpv1 labeling in the dorsal horn was unaltered in the PAM VGLUT2 cDKO^Trpv1^ mice (fig. S10). Altogether, this demonstrates that glutamate and neuropeptides function as parallel nociceptor neurotransmitter systems that are each sufficient to trigger a wide range of pain responses.

**Figure 5.**
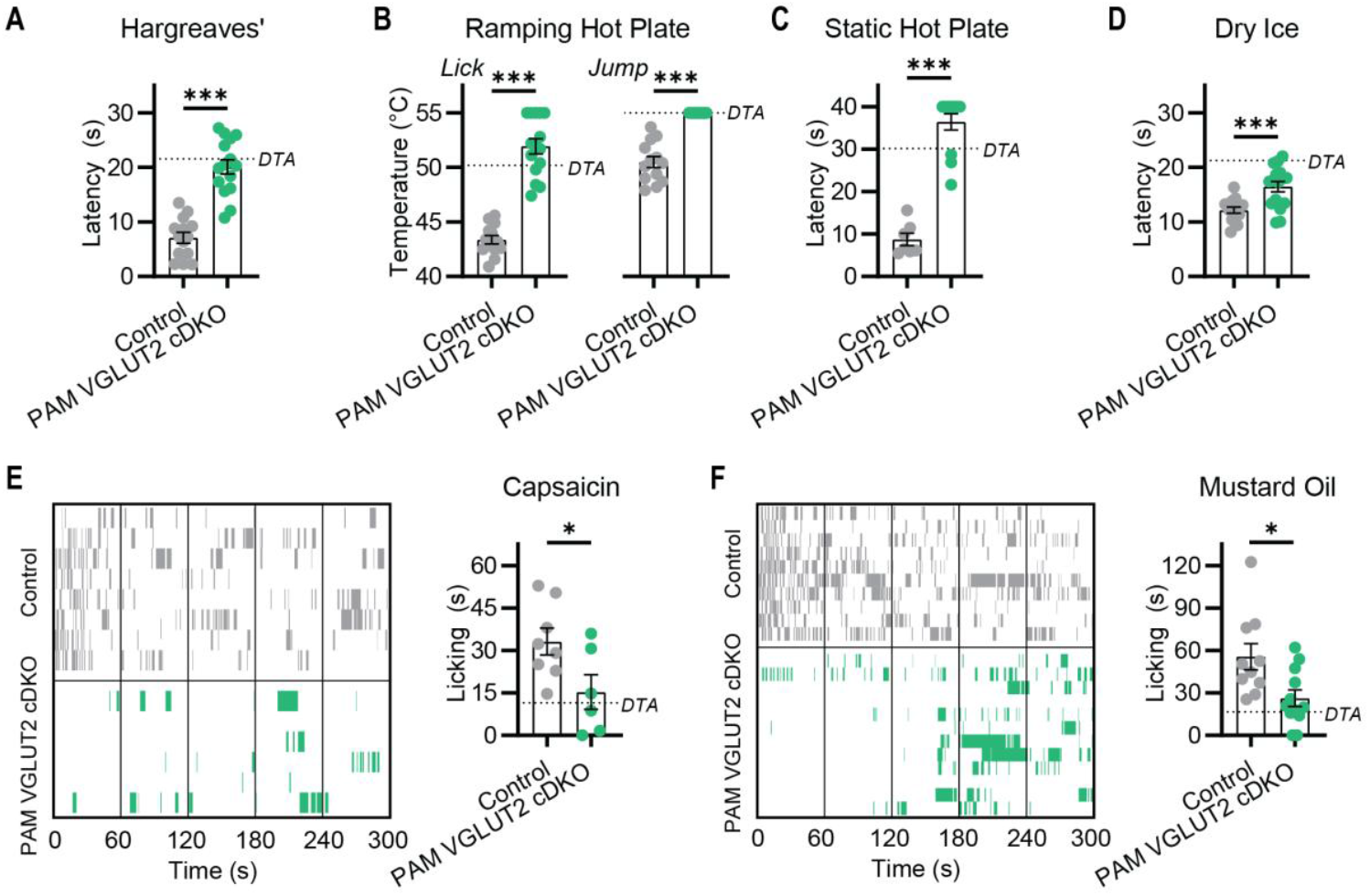
Mice lacking both PAM and VGLUT2 in nociceptors are insensitive to pain. **(A-F)** Pain behavioral assessment of PAM VGLUT2 cDKO mice using the same assays used for the single KO lines reveals these mice are profoundly insensitive to multiple types of acute painful stimuli. The dashed line denotes the mean values measured in DTA mice in Fig. 1. **(A)** Latency to withdrawal to radiant heat. n=14 (9M/5F) for Control and n=15 (8M/7F) for PAM VGLUT2 cDKO. **(B)** Temperature of first lick and jump on Ramping Hot Plate. n=13 (8M/5F) for Control and n=15 (8M/7F) for PAM VGLUT2 cDKO. **(C)** Latency to withdrawal on Static Hot Plate. n=6 (4M/2F) for Control and n=12 (8M/4F) for PAM VGLUT2 cDKO. **(D)** Latency to withdrawal to dry ice. n=14 (9M/5F) for Control and n=16 (11M/5F) for PAM VGLUT2 cDKO. **(E)** Ethogram and summary data showing time spent licking paw injected with capsaicin. n=8 (5M/3F) for Control and n=6 (5M/1F) for PAM VGLUT2 cDKO. **(F)** Ethogram and summary data showing time spent licking paw injected with mustard oil. n=10 (5M/5F) for Control and n=12 (6M/6F) for PAM VGLUT2 cDKO. Bars represent mean ± SEM. Means compared using an unpaired t test. *p<0.05, **p<0.01 ***p<0.001.

## Discussion

Since the discovery that nociceptors express and release Substance P (*37*), there has been heated debate about the role of neuropeptides in pain (*7, 38*). Though these neurons also release glutamate, we found that the effects of genetically inhibiting either transmitter system are nuanced and fail to recapitulate the loss of pain observed when nociceptors are ablated. By contrast, combined inhibition of both types of neurotransmitters caused profound analgesia, thus revealing how glutamate and neuropeptides can function independently to drive pain.

Disrupting the function of nociceptors in humans results in a congenital insensitivity to pain syndrome that leads to devastating injuries (*1*). Animals with corresponding mutations also lack pain responses and often exhibit unexplained wounds (*39*). The ability to detect potential or ongoing tissue damage is therefore critical for survival. Anecdotally, mice with Trpv1-Cre mediated deletion of both PAM and VGLUT2 showed more severe self-mutilation than either deletion alone. Intriguingly, this was never observed in the corresponding nociceptor ablated mice, likely showing that residual nociceptor signals (e.g., through the itch related NP3-cells that express the neuropeptides SST, NPPB and NTS, which are not C-terminally amidated) drive this phenotype (*28*). This highlights how a range of pain and itch transmitters may collectively choreograph the behavioral response to peripheral sensory inputs.

Our work shows that two very different neurotransmitter systems can evoke the same behavioral output, arguing that multiple pathways generate and converge on a shared pain-state in the spinal cord, reminiscent of mechanisms operating in the mammalian hypothalamus (*40, 41*) and lobster stomatogastric ganglion (*42*). Despite some notable successes (*6, 43, 44*), this type of degeneracy is a possible explanation for the failure of many novel targets to provide lasting pain relief (*45*–*47*). Our findings thus suggest that any perturbation of the spinal cord circuitry that prevents the common pain-state from emerging would reduce or even block pain therapeutically.

## Acknowledgments

We thank Dr Nick Ryba and members of the Chesler, Ryba and Hoon Labs for help and advice. We are grateful to Dr Lee Eiden for sharing the PACAP-flox mice, and to Dr Chuan Wu, Dr Hao Jin and their labs for help and guidance with pilot experiments. This work was supported by the National Center for Complementary and Integrative Health and National Institute of Neurological Disorders and Stroke (ZIA AT000028, ATC). DIM was supported by a European Molecular Biology Organization Postdoctoral Fellowship (ALTF 333-2021), Branco Weiss Fellowship – Society in Science, and the NIH Office of Autoimmune Disease Research Intramural Award. Bulk RNA Sequencing was performed by the NIDCR Genomics and Computational Biology Core.

## Disclaimer

This research was supported in part by the Intramural Research Program of the National Institutes of Health (NIH). The contributions of the NIH authors were made as part of their official duties as NIH federal employees, are in compliance with agency policy requirements, and are considered Works of the United States Government. However, the findings and conclusions presented in this paper are those of the authors and do not necessarily reflect the views of the NIH or the U.S. Department of Health and Human Services.

## Author contributions

Conceptualization: DIM, ATC

Methodology: DIM, RB, ATC

Investigation: DIM, RB

Visualization: DIM, RB

Funding acquisition: DIM, ATC

Project administration: ATC

Supervision: DIM, ATC

Writing – original draft: DIM, RB, ATC

Writing – review & editing: DIM, RB, ATC

## Competing interests

Authors declare that they have no competing interests.

## Data and materials availability

All data are available in the main text or the supplementary materials.

## Materials and Methods

### Animals

Animal care and experimental procedures followed protocols approved by the National Institute for Neurological Disorders and Stroke (NINDS) Animal Care and Use Committee. Trpv1-Cre (Jax #017769), VGLUT2-flox mice (Jax #012898), ROSA-LSL-DTA (Jax #009669), and Avil-Cre (Jax #032536) were purchased from Jackson Lab. PAM-flox mice were generated in-house as described in Ref. *(28)*. Hoxb8-Cre mice were a gift from Hanns Ulrich Zeilhofer (29) and PACAP-flox mice were a gift from Lee Eiden (*48*). Both male and female mice older than six weeks were included in all experiments, and the number of animals of each sex for each dataset is specified in the corresponding figure legend. All behavioral experiments were conducted with the experimenter blinded to genotype. Genomic DNA for genotyping was extracted from tail biopsies and analyzed by Transnetyx.

### Behavioral assays

#### Hargreaves

Mice were habituated in acrylic chambers on a plexiglass stand for 1 hour the day of and day before testing, with the Hargreaves machine (IITC) turned on in a dark room. The radiant heat stimulus was aimed for the centre of the plantar region of the mouse’s hind paw. The active intensity of the stimulus was 40 intensity units. Response latency was recorded for five trials with 10 min breaks in between each trial, and a median calculated. The cut-off time was 30 s.

#### Ramping Hot Plate

The Hot Plate (Ugo Basile) was set to the starting temperature of 25 °C. A mirror was placed behind the setup for better visibility of the mouse on the Hot Plate, and mice were recorded throughout the experiment using a web-camera (Logitech). A clear cylindrical tube was placed around the Hot Plate to ensure the mouse could not escape. Mice were placed on the Hot Plate and the top of the cylinder was covered to prevent escape. The mouse was allowed to explore the plate for 1 min at 25 °C, before the temperature was increased at a rate of +1 °C every 7 s in a pre-programed ramp to 55 °C. Once the temperature reached 55 °C, mice were immediately removed and returned to their home cage. The temperature at which a mouse first licked and jumped was determined by a blinded observer post-hoc using the recorded videos. The plate was thoroughly cleaned between mice.

#### Static Hot Plate

Mice were habituated and recorded as described in the preceding section. The Hot Plate (Bioseb) was maintained at a temperature of 50 °C based on a surface thermode reading. A rectangular chamber was placed around the hot plate and the top was covered to prevent escape. Mice were placed on the Hot Plate and observed carefully. The mice were immediately removed and returned to the home-cage following a positive response (licking paws or jumping), or when a cut-off time of 40 s was reached. The precise latency to lick was scored by a blinded observer post-hoc using the recorded videos.

#### Dry ice

Mice were habituated in acrylic chambers on a plexiglass platform for 1 hour the day of and day before testing. A 2 ml plastic syringe was cut in half to allow for crushed dry ice to be compacted into the tube. The dry ice was pushed up against the glass where the mouse’s hind paw was resting. Withdrawal latency was recorded with a stopwatch. 10 minutes were given in between each trial. The procedure was repeated five times, and a median of the withdrawal latencies was calculated.

#### Chemical algogens

Mice were habituated in acrylic chambers on a plexiglass platform (capsaicin) or wire mesh (mustard oil) for 1 hour the day of and day before testing. 20 µl of 0.3 mM capsaicin in 90% saline / 9% Tween-80 / 1% EtOH, or 30 µl of 1% AITC in saline was injected into the plantar surface of the hind paw. Mice were recorded for 5 minutes following algogen injection. Using BORIS software, a blinded observer scored individual bouts of paw-directed licking as state events over the 5 minute time period. The overall time spent licking was calculated using the Time Budget function, and ethograms were generated using the Plot function. Cumulative licking time for all recorded events from each genotype was plotted against time, and normalized to the number of mice.

### Neuropeptide imaging

#### Generation and maintenance of biosensor cell lines

The generation and characterization of NKR1 and CLR biosensors is described in Ref. *(18)*. To generate the new biosensor cell lines, we produced neuropeptide receptor DNA constructs for Tacr2, Nmbr, Galr1, Adcyap1r1, Sstr2, Ntsr1, Oprd1, and Grpr encoding NKR1, NMBR, GALR1, PACR1, SSTR2, NTSR1, DOR and GRPR respectively (Epoch Life Science). Each receptor was subcloned along with a synthesized human G-protein α-subunit gene Gα15 and GCaMP6s into a dox-inducible backbone (GS66685-1). These were transfected into the Flp-In T-REx HEK293 cell line (Thermo Fisher Scientific, R78007) using a PiggyBac transposase system and stably-transfected cells were isolated using puromycin (1-10 μg/ml) to generate a polyclonal cell line. Doxycycline (2-5 μg/ml) was then used to induce expression of the construct.

The cell lines were maintained on polystyrene culture plates (Thermo Fisher Scientific, 07-200-80) in a 5% CO2 humidified incubator at 37 °C. The growth medium was changed every 2–3 days and consisted of DMEM/F12 (Thermo Fisher Scientific, 11330032) supplemented with 10% fetal bovine serum (Thermo Fisher Scientific, 26140079) and puromycin. Cells were passaged when they reached confluence, which was roughly twice per week, and were never propagated beyond 20 passages. For passaging, cells were rinsed in PBS (Thermo Fisher Scientific, 10010023) and then incubated in Accutase (Thermo Fisher Scientific, 00-4555-56) for ∼5 minutes at 37 °C to detach. Cells were collected in a 15 ml tube (Thermo Fisher Scientific, 12-565-268) and centrifuged at 300 rcf for 3 minutes to pellet. The supernatant was aspirated, and cells were resuspended in growth medium followed by plating in new polystyrene plates. Typical dilution ratios for passaging were between 1:5 and 1:20. For the initial characterization, biosensor cell lines were cultured onto eight-well glass slides with approximately 100,000 cells per well, in the presence of doxycycline and imaged when they reached confluency.

#### Adult dorsal root ganglion culture and biosensor cell co-culture

Dorsal root ganglia (DRG) were dissected from the entire length of the vertebral column and then digested in a pre-equilibrated enzyme mix for 35–45 min (37 °C, 5% CO2). The enzyme mix consisted of Hanks’ balanced salt solution containing collagenase (type XI; 5 mg/ml), dispase (10 mg/ml), HEPES (5 mM), and glucose (10 mM). DRGs were then gently centrifuged for 3 min at 300 revolutions per minute, the supernatant was discarded and replaced with 1 ml of warmed DMEM/F-12, supplemented with 10% fetal bovine serum (FBS). Next, DRGs were mechanically triturated approximately 25 times using a 1 ml pipette. 1 ml of the same media containing 15% Bovine serum albumin solution (BSA) was then added to the dissociated cells to produce a density gradient. Dissociated cells were then centrifuged again at 300 revolutions per minute, the supernatant was discarded and cells were re-suspended in the required volume of DMEM supplemented with FBS, and a growth factor cocktail consisting of nerve growth factor (mouse beta-NGF, 25 ng/ml), neurotrophin-3 (human NT-3, 25 ng/ml), brain-derived neurotrophic factor (human/mouse/rat BDNF, 25 ng/ml), glial cell line-derived neurotrophic factor (mouse GDNF, 25 ng/ml). Finally, cells were plated onto 8-well glass slides previously coated with poly-L-lysine (1 mg/ml) and laminin (1 mg/ml), and incubated at 37 °C in 5% CO2. 1 hour later the wells were filled with media containing AAV9-CAG-tdTomato (codon diversified) (Addgene, 59462-AAV9, 1:200 dilution). After 72 hours, DRG neurons had grown dense processes and could be identified by tdTomato fluorescence. At this 72 hr time point, biosensor cells were resuspended in DMEM/F-12, supplemented with 10% fetal bovine serum (FBS), the growth factor cocktail and doxycycline (2-5 μg/ml). Between 150-200,000 cells were dispensed into each imaging well containing DRG neurons, and incubated for at least a further 24 hr before imaging was performed at 96 hr.

#### In vitro imaging

Imaging was performed in Ringer’s solution: 125 mM NaCl, 3 mM KCl, 5 mM CaCl2, 1 mM MgCl2, 10 mM glucose, and 10 mM HEPES (all from Sigma-Aldrich), adjusted to pH 7.3 with 1 M NaOH, and osmolality measured ∼280 mmol/kg. Biosensor cells alone, or co-cultured with DRGs, were rinsed in Ringer’s solution and imaged in Ringer’s solution at room temperature on an Olympus IX73 inverted microscope using a 10x objective and pco.panda sCMOS back-illuminated camera at 1 frames/s. tdTomato was visualized using a 560 nm LED and GCaMP6f at 488 nm. All imaging trials began with 15 s of baseline measurement and then the cells were treated with chemicals by micropipette. The chemicals used included capsaicin (10 μM), and a range of neuropeptides (1 pM to 10 μM): Substance P, CGRP, Neurokinin A, Neuromedin B, Galanin, PACAP1-38, Somatostatin, Neurotensin, Leu-enkephalin and Gastrin-releasing peptide.

Fiji software was used to import and analyze video files from pco.camware software (pco). To remove application artefacts, the background was reduced using the background subtraction function with a rolling ball radius of 50 pixels. The peak ΔF/F_0_ across the entire field of view was calculated to quantify the changes in fluorescence associated with chemical application, with F_0_ defined as the mean fluorescence intensity of the entire field of view in the 5 s immediately preceding stimulation. The ΔF/F_0_ over the 75 s post application was used to calculate area under the curve to provide an integrated measure of receptor activation.

### Immunohistochemistry

Under 2% isoflurane anesthesia, mice were perfused intracardially with 0.01% heparin in PBS followed by 4% PFA in PBS. DRG and spinal cords were then dissected and stored in 4% PFA solution overnight. Following overnight fixation, the tissue was cryoprotected in 30% sucrose solution in PBS containing 0.01% Azide. Tissue was then mounted using OTC and cut into 30-40 µm sections using a cryostat. Sections were immediately stored in 0.01% Azide solution in PBS.

Immunohistochemistry began with incubating sections in a blocking buffer solution (5% normal donkey serum; 0.1% Triton X-100 in PBS) on a shaker at room temperature for 3 hours. Afterward, sections were incubated in blocking solution containing a subset of the following primary antibodies: 1:500 or 1.1000 goat anti-CGRP polyclonal primary antibody (Abcam, #ab36001), 1:500 rat anti-Substance P polyclonal primary antibody (Abcam, #ab67006), 1:500 rabbit anti-TRPV1 polyclonal primary antibody (Alomone Labs, #ACC-030), 1:500 chicken anti-Peripherin polyclonal primary antibody (Invitrogen, #PA1-10012), 1:1000 guinea pig anti-VGlut2 polyclonal primary antibody (Millpore Sigma, #AB2251-I), and 1:1500 rabbit anti-PAM polyclonal primary antibody (Proteintech, #26972) on a shaker at room temperature overnight.

The following day, the tissue sections were rinsed 4 times with PBS and then incubated in blocking solution containing a subset of corresponding secondary antibodies: 1:200 RRX-conjugated donkey anti-rabbit secondary antibody, 1:200 Cy5-conjugated donkey anti-rat secondary antibody, 1:200 FITC-conjugated donkey anti-goat secondary antibody, 1:200 FITC-conjugated donkey anti-chicken secondary antibody, 1:200 FITC-conjugated donkey anti-guinea pig secondary antibody, 1:200 Cy5-conjugated donkey anti-rabbit secondary antibody, 1:200 FITC-conjugated donkey anti-rat secondary antibody (Thermo Fisher Scientific), or 1:200 750-conjugated donkey anti-goat secondary antibody (Abcam, #AB175744).

After rinsing sections 4 times in PBS, the sections were mounted onto glass slides (Daigger Scientific) and coated with ProLong diamond antifade mounting media (Thermo Fisher Scientific). Sections were imaged in Z stacks with an Olympus confocal microscope under a 10x objective lens.

In one experiment, Alex Fluor 555-conjugtated cholera toxin-B (10 μl, 0.1% in PBS) was injected into the mouse hindpaw 7 days before tissue harvest, to enable visualization of paw-derived axon terminals in the dorsal horn of the spinal cord.

### Laser speckle contrast imaging (LSCI)

Mice were anesthetized using isofluorane (2-3%) in a box. Once clearly anesthetized, they were moved to a nose cone, secured with incisor bars, and maintained with anesthetic at approximately 1.5% at a rate of around 500 ml / min on a warming pad. The pedal reflex was used to assess the stability of anesthesia. Once stable, paws were immobilized using double-sided tape attached to dorsal side, such that the plantar surface of the paw faced up. Dermal blood flow in the hindpaw was then measured by laser speckle contrast imaging (moorFLPI-2, Moor Instruments, Wilmington, DE, USA) by monitoring flux in the imaging window across time. The LSCI video was continuously acquired at 1 Hz. The spatial processing mode was used, with a high-resolution sliding window and a low pass filter with a 1 s time constant. Following at least a 1 min stable baseline, 10 μl of 5% mustard oil was applied to the paw to activate nociceptor terminals, and the recording continued for 5 mins to monitor changes in dermal blood flow. Greyscale images were also collected for use as example images and for quality control.

LSCI videos were exported from the Moor instruments software as AVI files. The AVI files were opened in FIJI. To remove any lateral movement, videos were aligned using the turboreg plugin (rigid body). A region of interest was drawn around the paw using the wand tool. The mean flux time course was exported. To quantify the changes in flux associated with mustard oil application, Δflux was calculated using the procedure used to calculate ΔF/F_0_, with baseline (F_0_) defined as the mean flux of the region of interest in the 15 s immediately preceding mustard oil stimulation. The Δflux was integrated over time to calculate an area under curve. This was calculated from 15 s post stimulation to prevent the application artefact contaminating the signal.

### Bulk RNA sequencing

To conduct bulk RNA sequencing of sensory neurons in adult PAM WT (n = 2 female, n = 2 male) and PAM cKO mice (n = 2 female, n = 2 male), DRGs (14-15 per animal) were first dissected and placed into DNA/RNA Shield™ stabilization solution (Quick-RNA™ Miniprep Kit, Zymo Research R1054). Afterward, the tissue was mechanically homogenized using the IKA T 10 Basic ULTRA-TURRAX® dispersing instrument. Immediately after homogenization, each sample tube was placed on dry ice and then moved to a −80°C freezer. After thawing the homogenized DRG samples to room temperature, total RNA was purified using the Quick-RNA™ Miniprep Kit (Zymo Research, Irvine, CA) according to manufacturer’s instructions. RNA yields were measured using the Qubit instrument and ranged from 70-130 ng/µL.

Bulk RNA sequencing was performed by the NIDCR Genomics and Computational Biology Core. Mouse DRG libraries were produced by the AZENTA/GENEWIZ using a Nextera/Proprietary method. Samples were run on an Illumina NovaSeq6000 configured for 150 PE reads. FASTQ files were extracted using bcl2fastq, and were further processed using a standard analysis pipeline to determine read quality, sequencing saturation and differential gene expression. Three independent statistical tests were used to assess differential gene expression between groups: DeSeq2, edgeR and Limma-Voom. All three produced similar results with only minor changes in gene expression between control and PAM cKO samples. The differential gene expression analysis using DeSeq2 is shown in fig. S6.

### Analysis of gene expression

Single-cell sequencing data from DRG (*49*) were obtained from GEO Series GSE254789 and analysed with Seurat v.5 in RStudio. Cells with less than 800 expressed genes or more than 5% of mitochondrial transcripts were excluded, and datasets were combined using canonical correlation analysis integration after principal component analysis reduction to 30 components. Neuronal and non-neuronal cell clusters were identified in UMAP by analyzing the expression of Snap25, Mbp, Apoe, Qk, Pecam1, Slc17a7 and Slc17a6. Doublets were identified using DoubletFinder v.3, and doublets and non-neuronal cells were removed from the dataset. After neurons were renormalized, reduced to 40 principal components and reintegrated, neuronal clusters were calculated using the Louvain algorithm with a resolution of 0.2 and identified/combined on the basis of the marker genes used in Ref. (*50*).

### Statistics

Data were compared using two-way ANOVA with post-hoc Hom-Sidak’s test, One-WAY ANOVA with post-hoc Holm-Sidak’s test, and Student’s unpaired t test. The α value was 0.05. For all experiments n is the number of animals, except biosensor imaging where n is the number of wells. Error bars denote standard error of the mean throughout. No power analyses were used to determine sample sizes, but our sample sizes are similar to those from previous studies (12, 18, 19). Graphs were generated and statistical analysis was performed using GraphPad 8.0 software (Prism). All statistical tests and results are reported in the figure legends.

## Supplementary Figures

**Figure S1.**
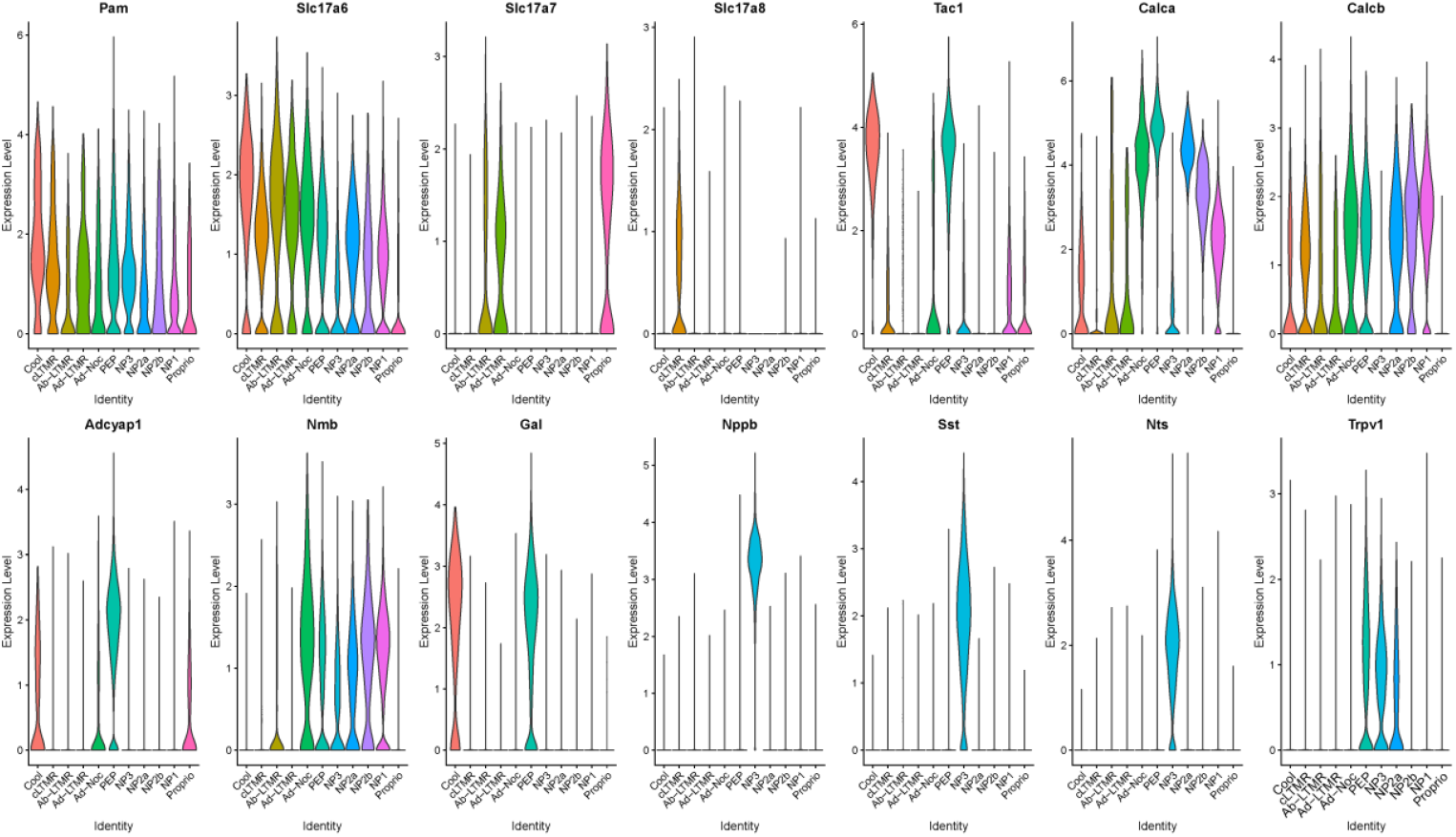
Subset-specific expression of neurotransmitter and neuropeptide-related genes in mouse dorsal root ganglia neuron subtypes. Violin-plot analysis of expression level (log normalized single cell RNA sequencing data) *(49)* of neuropeptide- and neurotransmitter-related genes across the 10 sensory neuron transcriptomic classes used in Ref. *(50)*.

**Figure S2.**
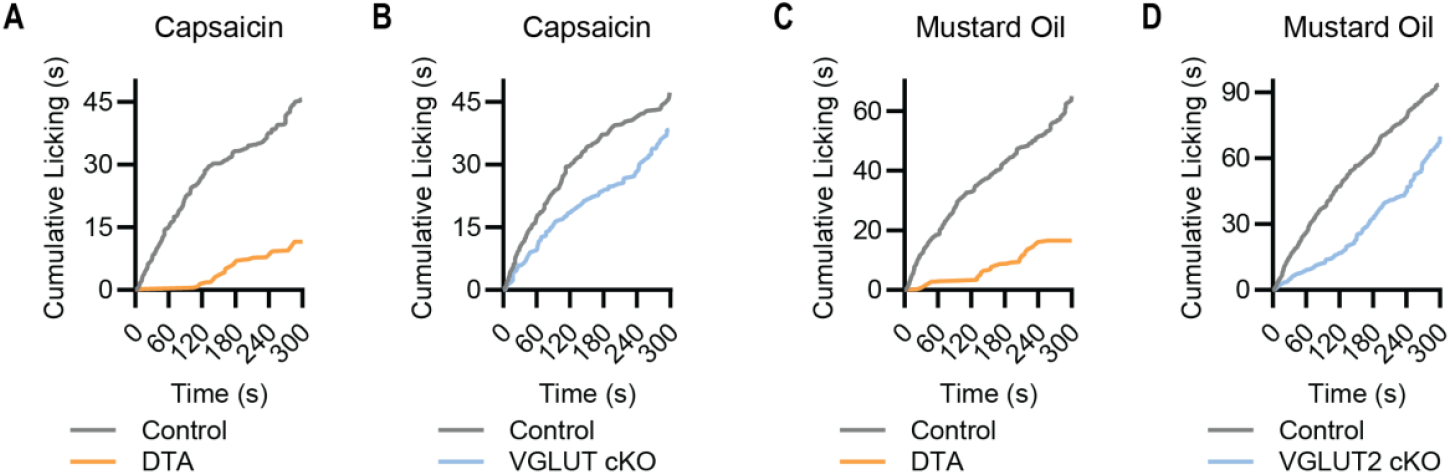
Temporal structure of chemical pain in DTA and VGLUT2 cKO mice. **(A-D)** Cumulative paw licking (normalized to number of mice per group) following capsaicin or mustard oil injection in DTA mice, VGLUT2 cKO mice, and their respective littermate controls. For capsaicin, (A) n=9 (5M/4F) for Control and n=5 (4M/1F) for DTA; (B) n=5 (3M/2F) for Control and n=7 (5M/2F) for VGLUT2 cKO. For mustard oil, (C) n=9 (5M/4F) for Control and n=5 (4M/1F) for DTA; (D) n=10 (3M/7F) for Control and n=12 (5M/7F) for VGLUT2 cKO.

**Figure S3.**
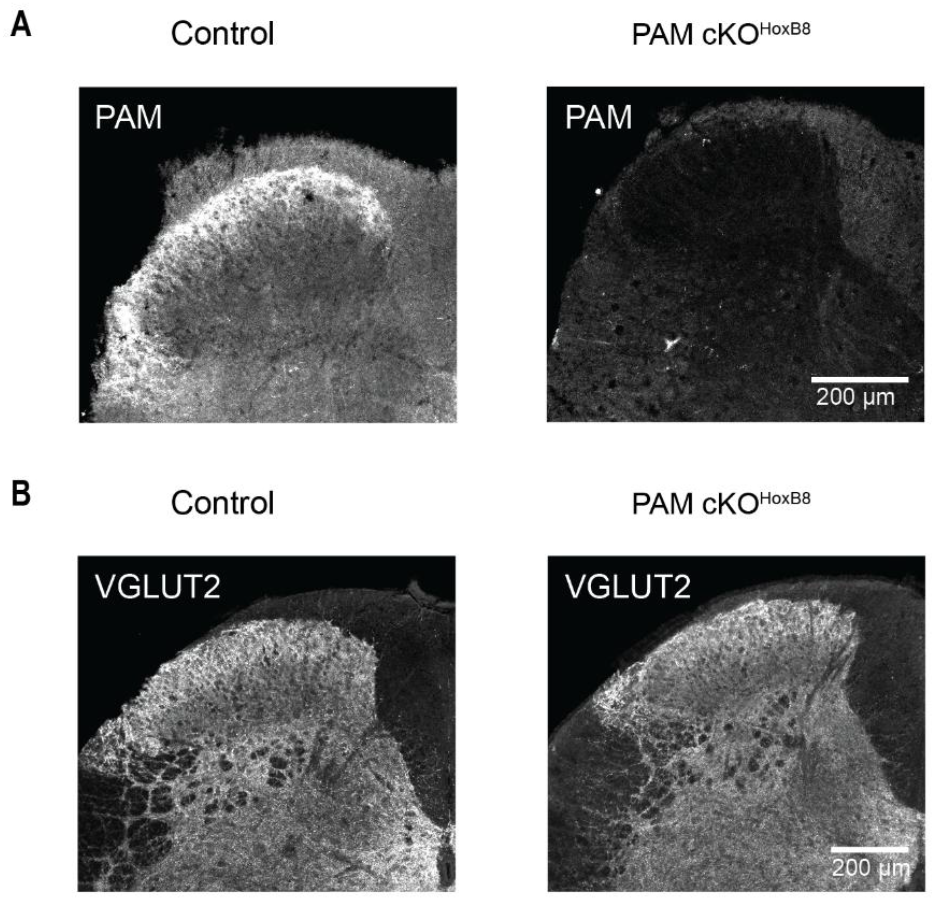
Immunohistochemical staining of PAM cKO^Hoxb8^ dorsal horn. **(A)** Example confocal image showing that PAM is enriched in the superficial laminae of the dorsal horn of control mice (left). This staining is completely absent in the PAM cKO_Hoxb8_ (right). Note the overall reduction in staining intensity due to PAM deletion in all spinal cord cells in this mouse. **(B)** Example confocal image showing that VGLUT2 staining is similar in control and PAM cKO_Hoxb8_ dorsal horn.

**Figure S4.**
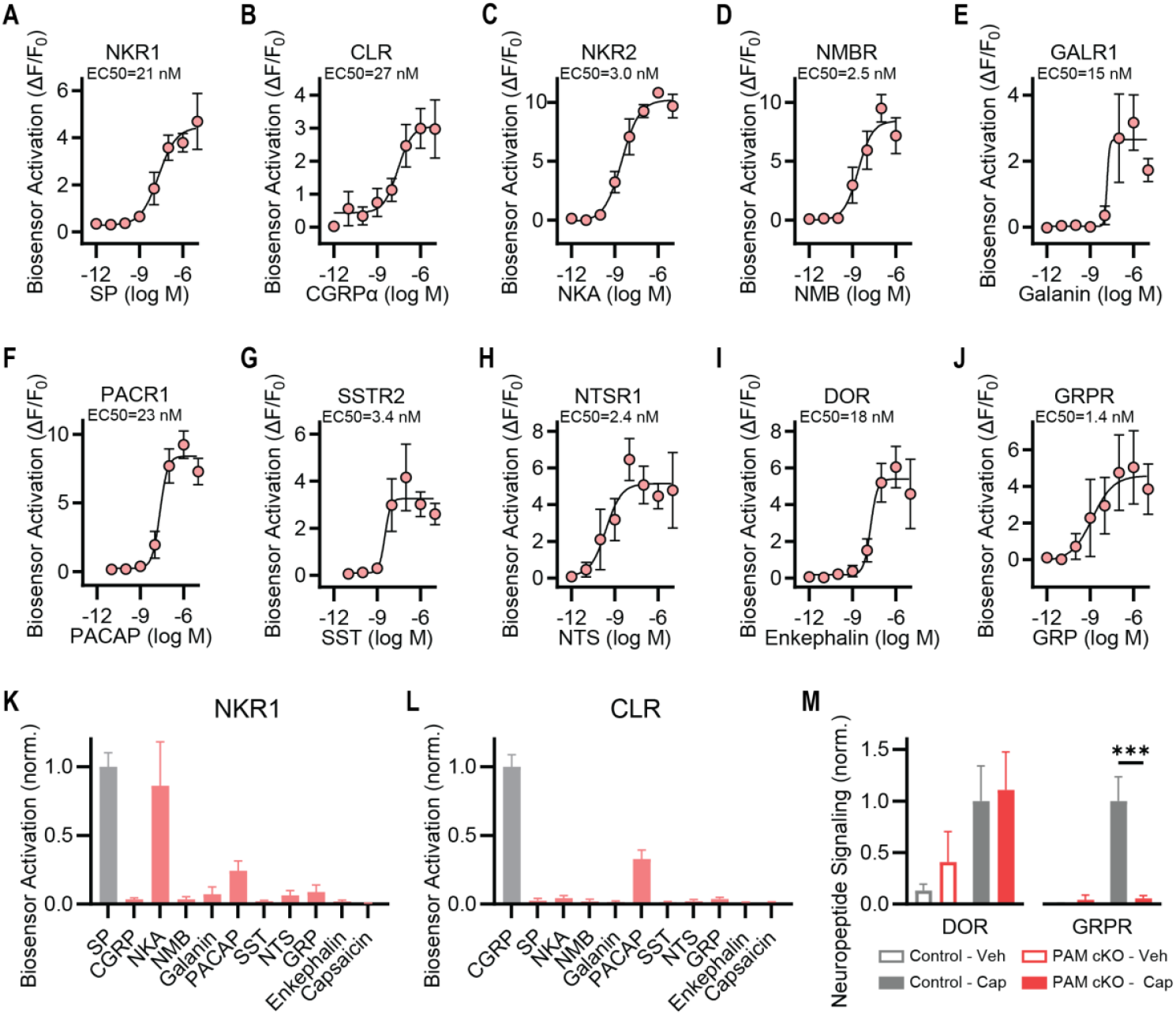
Characterization of neuropeptide biosensors. **(A-J)** Dose response curves showing the sensitivity of each GPCR biosensor cell line to its target neuropeptide. Four-parameter variable slope dose-response curves were fit by non-linear regression. At least three replicates were acquired for each concentration, in two independent experiments. Please note the NKR1 and CLR data in (A) and (B) were previously published in Ref. (*18*), and are shown for comparison with the new cell lines. **(K)** Response of NKR1 cell line to a panel of neuropeptides. As expected, these cells are only activated by the tachykinins SP and NKA. **(L)** CLR cell lines specifically respond to SP. For (K) and (L), all neuropeptides were at 1 μM and capsaicin was at 10 μM. **(M)** Signaling by Enkephalin and GRP can be detected using DOR and GRPR biosensors from control nociceptors following capsaicin stimulation. PAM cKO nociceptors evoke normal DOR activity, but there is a loss of GRPR signaling. DOR: n=9 for Control; n=7 for PAM cKO. GRPR: n=10 for Control; n=13 for PAM cKO. Bars represent mean ± SEM. For each biosensor, means were compared using 2-way ANOVA followed by post-hoc Holm-Sidak’s multiple comparison’s test. *p<0.05, **p<0.01 ***p<0.001.

**Figure S5.**
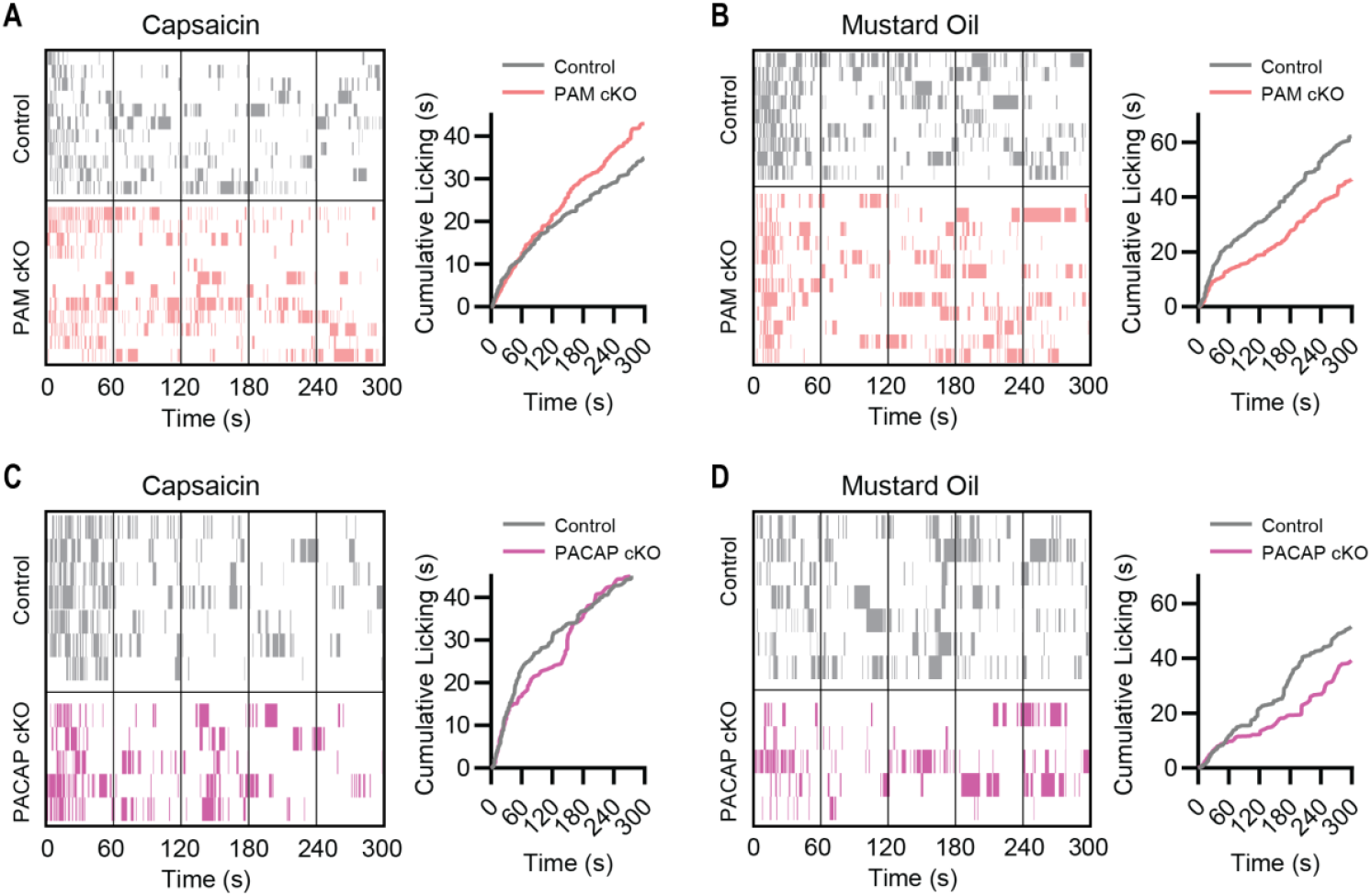
Temporal structure of chemical pain in PAM and PACAP cKO mice. **(A-D)** Ethograms and normalized cumulative licking plots for littermate control, PAM cKO and PACAP cKO mice given intraplantar injections of the chemical algogens capsaicin and mustard oil. (A) For capsaicin, n=12 (6M/6F) for Control and n=12 (8M/4F) for PAM cKO. (B) For mustard oil, n=9 (2M/7F) for Control and n=12 (3M/9F) for PAM cKO. (C) For capsaicin, n=7 (4M/3F) for Control and n=5 (3M/2F) for PACAP cKO. (D) For mustard oil, n=7 (4M/3F) for Control and n=5 (3M/2F) for PACAP cKO.

**Figure S6.**
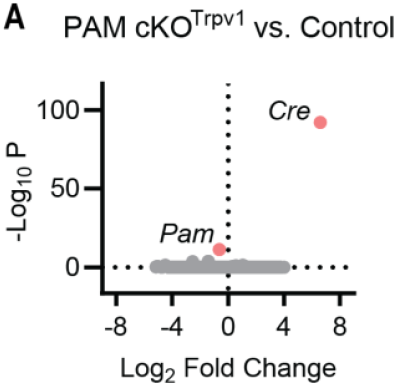
Deletion of PAM in Trpv1 afferents does not cause marked transcriptomic changes. **(A)** Bulk RNA sequencing shows very few genes are differentially expressed between control and PAM cKO dorsal root ganglia. As expected, Cre was enriched in PAM cKO_Trpv1_ DRGs, and PAM was downregulated. n=4 (2M/2F) for Control; n=4 (2M/2F) for PAM cKO_Trpv1_.

**Figure S7.**
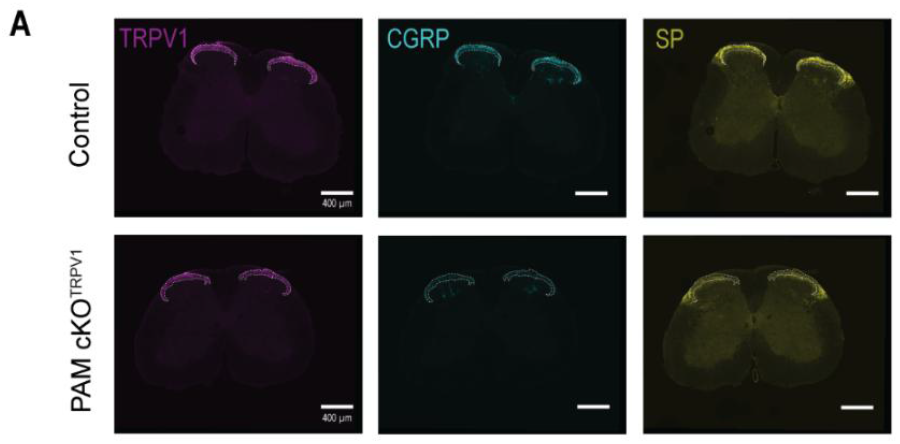
Deletion of PAM in Trpv1 afferents reduces neuropeptide staining. **(A)** Example confocal images of spinal cord dorsal horns showing that PAM deletion reduces the intensity of CGRP and Substance P (SP) staining in the terminal field of Trpv1 afferents. The dotted lines are traced from the region of Trpv1 staining. Note that only in the area of Trpv1 innervation would we expect to see reduced neuropeptide staining using the Trpv1-Cre approach.

**Figure S8.**
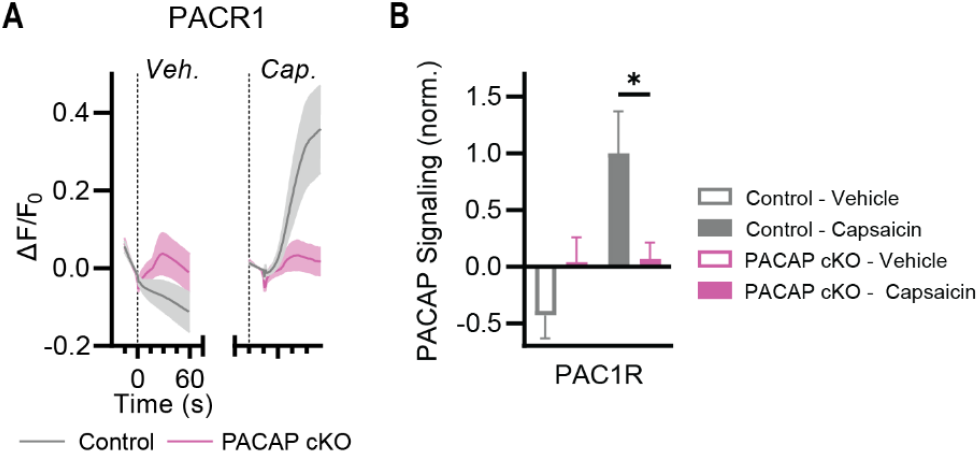
Loss of PACAP signaling from nociceptors in PACAP cKO mice. **(A)** Time course of PACAP release from nociceptors following capsaicin stimulation shows a complete loss in the PACAP cKO. **(B)** Summary plots showing PACAP signaling is abolished following PACAP deletion. n=13 from 2 Control mice; n=16 from 3 PACAP cKO mice. Bars represent mean ± SEM. Means were compared using 2-way ANOVA followed by post-hoc Holm-Sidak’s multiple comparison’s test. *p<0.05.

**Figure S9.**
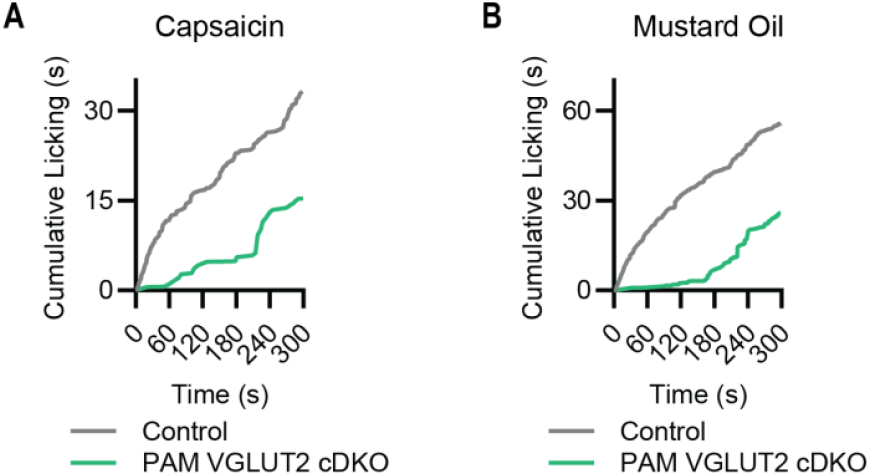
Temporal structure of chemical pain in PAM VGLUT2 cKO mice. **(A)** Cumulative licking (normalized to number of mice per group) to capsaicin. n=8 (5M/3F) for Control and n=6 (5M/1F) for PAM VGLUT2 cDKO. **(B)** Mustard oil-evoked cumulative licking. n=10 (5M/5F) for Control and n=12 (6M/6F) for PAM VGLUT2 cDKO.

**Figure S10.**
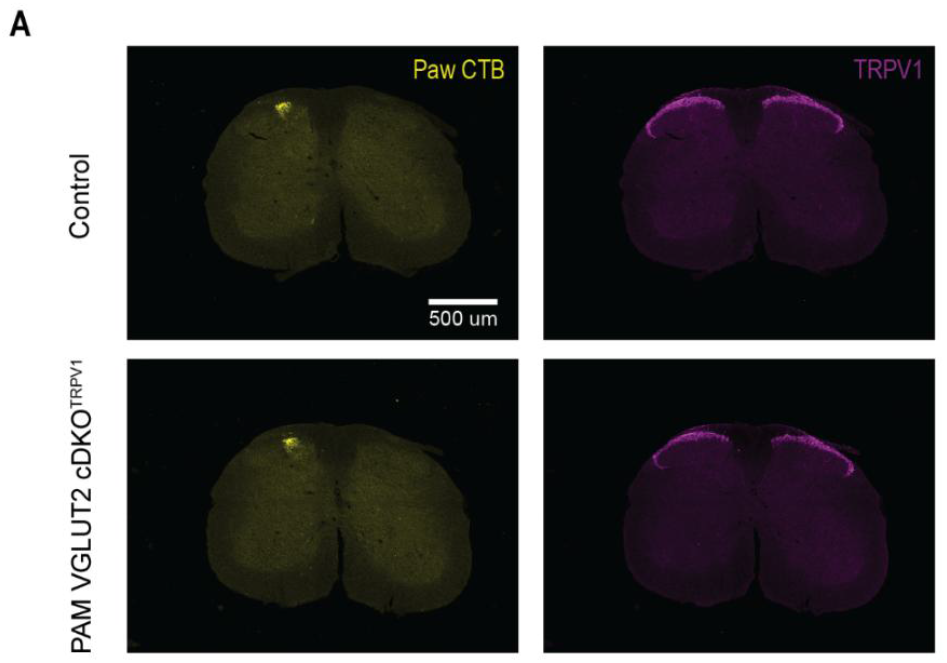
Combined deletion of PAM and VGLUT2 does not affect central terminal innervation. Example confocal images showing spinal cord dorsal horn staining for Trpv1 and cholera toxin B injected into the paw in Control (top) and PAM VGLUT2 cDKO mice (bottom).

## Notes

### Competing Interest Statement

The authors have declared no competing interest.

### Summary of Updates

We corrected the spelling of an authors name.

